# *Doublesex* mediates the development of sex-specific pheromone organs in *Bicyclus* butterflies via multiple mechanisms

**DOI:** 10.1101/686477

**Authors:** Anupama Prakash, Antónia Monteiro

## Abstract

The *Bicyclus* lineage of satyrid butterflies exhibits male-specific traits, the scent organ complex, used for chemical communication during courtship. This complex consists of tightly packed brush-like scales (hair-pencils) that rub against scent patches to disperse pheromones, but the evolution and molecular basis of the organ’s male-limited development remains unknown. Here, we examine the evolution of the number and location of the scent patches and hair-pencils within 53 species of *Bicyclus* butterflies, and the involvement of the sex determinant gene *doublesex (dsx)* in scent organ development in *Bicyclus anynana* using CRISPR/Cas9. We show that scent patches and hair-pencils arose via multiple, independent gains, in a correlated manner. Further, an initially non-sex-specific Dsx protein expression pattern in developing wing discs becomes male-specific and spatially refined to areas that develop the scent organ complex over the course of development. Functional perturbations of *dsx* show that this gene is required for male patch development whereas hair-pencils can develop in both sexes without Dsx input. Dsx in females is, instead, required to repress hair-pencils. These findings suggest that the patches and hair-pencils evolve as correlated composite organs that are sex-limited via the spatial regulation of *dsx*. Divergence in the function of *dsx* isoforms occurs in both sexes, where the male isoform promotes patch development in males and the female isoform represses hair-pencil development in females, both leading to the development of male-limited traits. Furthermore, evolution in number and location of patches, but not of hair-pencils, appears to be regulated by spatial regulation of *dsx.*

## Introduction

Diverse sex-specific traits are present in many animal lineages, arising as products of natural or sexual selection acting predominantly on one sex versus the other. Examples of such traits include variation in body size, color, exaggerated morphologies or behaviors and physiologies that aid each sex in acquiring more mates and/or in producing a larger number of offspring (Jonsson & Alerstam 1990; Gross 1996; Shine 2015). The enormous diversity and spectacular nature of some of these sex-specific traits combined with their important ecological and adaptive functions has promoted multiple studies looking at the evolution and development of such traits (Williams & Carroll 2009; Kopp 2012; Rogers *et al.* 2013; Gotoh *et al.* 2014; Neville *et al.* 2014).

The origin of sexually dimorphic traits is especially intriguing from a genetic perspective and was intensely debated by Darwin and Wallace (Kottler 1980). The two men debated whether novel traits can arise in a single sex from the very beginning or whether novel traits always arise first in both sexes but are then lost in one to create dimorphisms. In addition, each supported the importance of different selective forces in producing such dimorphisms. Wallace strongly believed that natural selection was key to converting equally inherited, showy traits to sex-limited traits for protection (primarily in females), while Darwin argued that traits could arise in only one sex (primarily in males) from the very beginning and be further amplified via sexual selection (Kottler 1980). This debate is only now being resolved with the use of sex-specific reconstructions of traits on phylogenies which help reconstruct single-sex or dual-sex origins of traits as well as their subsequent evolution (Emlen *et al.* 2005; Kunte 2008; Oliver *et al.* 2011). While it appears that both modes of evolution occur in a variety of taxa, the genetic and developmental mechanisms that allow the development of traits in one sex but limit their occurrence in the other sex, are still largely unresolved for most sexually dimorphic traits.

The scent organs in butterflies are a clear example of a sexually dimorphic, male-specific trait used for close-range, pre-mating chemical communication. These composite organs, collectively called androconia, differ dramatically in shape, color, pheromone composition and location, occurring as complexes of scent patches with modified epidermal scales and tightly-packed, brush-like hair-pencils on the legs, abdomen, thorax, and wings of butterflies that help produce and release pheromones during courtship (Boppre 1989; Birch *et al.* 1990; Doerge 2002; Hall & Harvey 2002; Hernández-roldán *et al.* 2014). In some species, some of the structures in the complex develop on different parts of the body and require special behaviors to ensure contact (Boppre 1989). For example, many male butterflies in the tribe Danaini (family Nymphalidae) have extrusible, brush-like abdominal hair-pencils that are brought into physical contact with pheromone producing patches on the wings to enable pheromone dissemination (Boppre 1989). On the other hand, many species in the tribe Satyrini possess scent organs consisting of hair-pencils (Fig 1 A,B) and scent patches (Fig 2A), both on the wing, that brush against each other dispersing chemicals in the process (Nieberding *et al.* 2008; Brattström *et al.* 2015, 2016; Aduse-Poku *et al.* 2017). It is to be noted however, that glandular, secretory cells underly some of the patches and produce pheromones, while other patches are not associated with such secretory cells (Bacquet *et al.* 2015; Dion *et al.* 2016).

**Figure 1:**
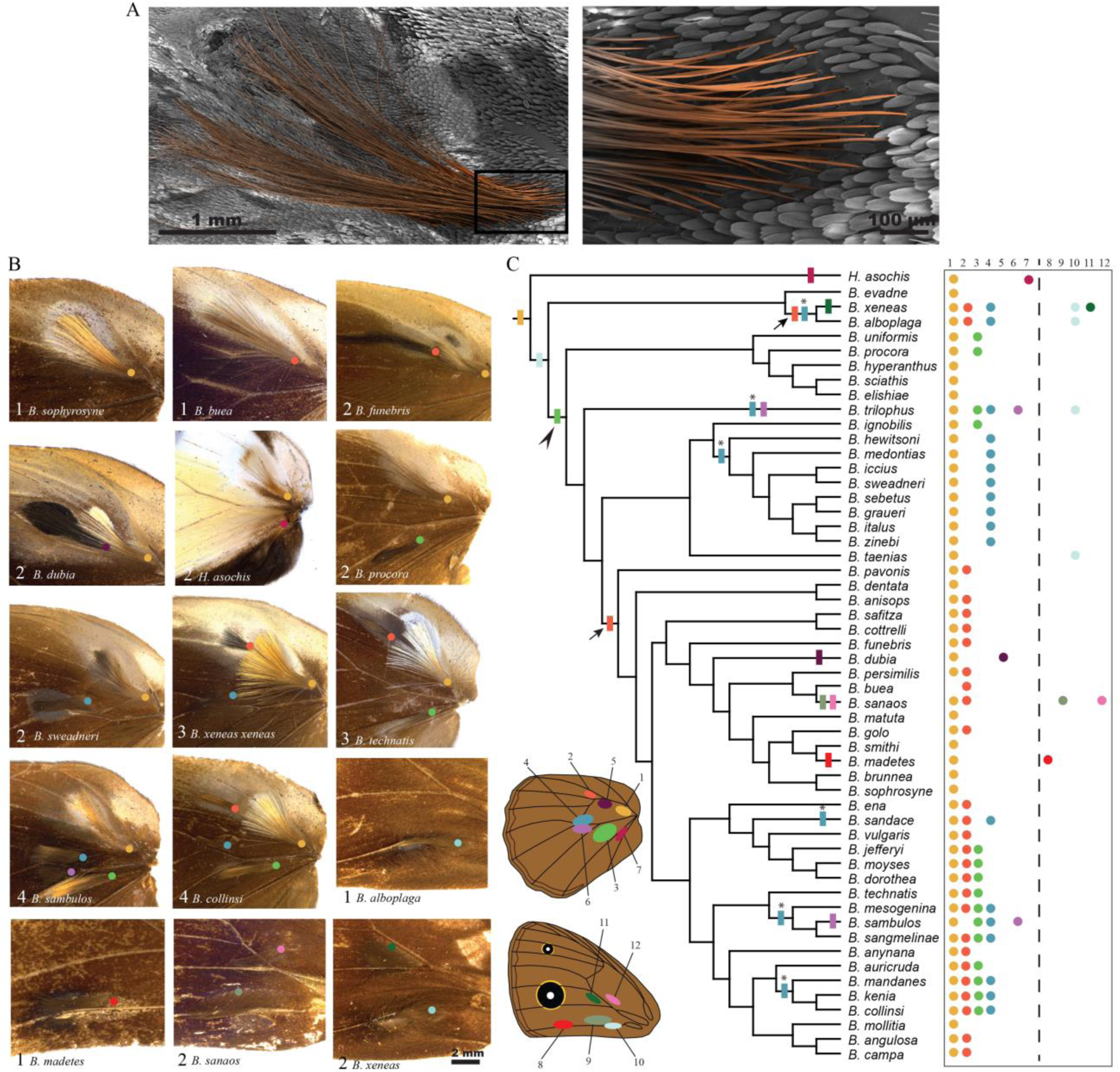
Diversity and evolution of hair-pencils on the wings of *Bicyclus* butterflies. A) Artificially colored scanning electron micrograph of the hindwing, yellow hair-pencil (hair-pencil 1) in *Bicyclus anynana*. The boxed area (base of the hair-pencil scales) is expanded to the right to illustrate the elongated and densely packed nature of the hair-likes scales that make up a hair-pencil. B) The different combinations of hindwing and forewing hair-pencils in male *Bicyclus* butterflies. The number of hair-pencils and the name of the species is denoted at the bottom left. Colored circles indicate the position of the base of each hair-pencil corresponding to the schematic in C. C) Evolutionary history of hair-pencils at homologous positions on the wing within the *Bicyclus* lineage. Locations, number and color codes of the different traits are given in the schematic and their presence/absence in each species is listed to the right. Filled rectangles on the phylogenetic tree indicate the likely gains of the traits for which either a single or a multiple-origin scenario at the MRCA of all the species bearing that trait was significantly supported. Arrows indicate the two likely gains for hair-pencil 2 and stars indicate the six likely gains for hair-pencil 4 within *Bicyclus*. For traits where a single- or multiple-origins scenario was equally supported, a single origin was mapped (black arrowhead).

**Figure 2:**
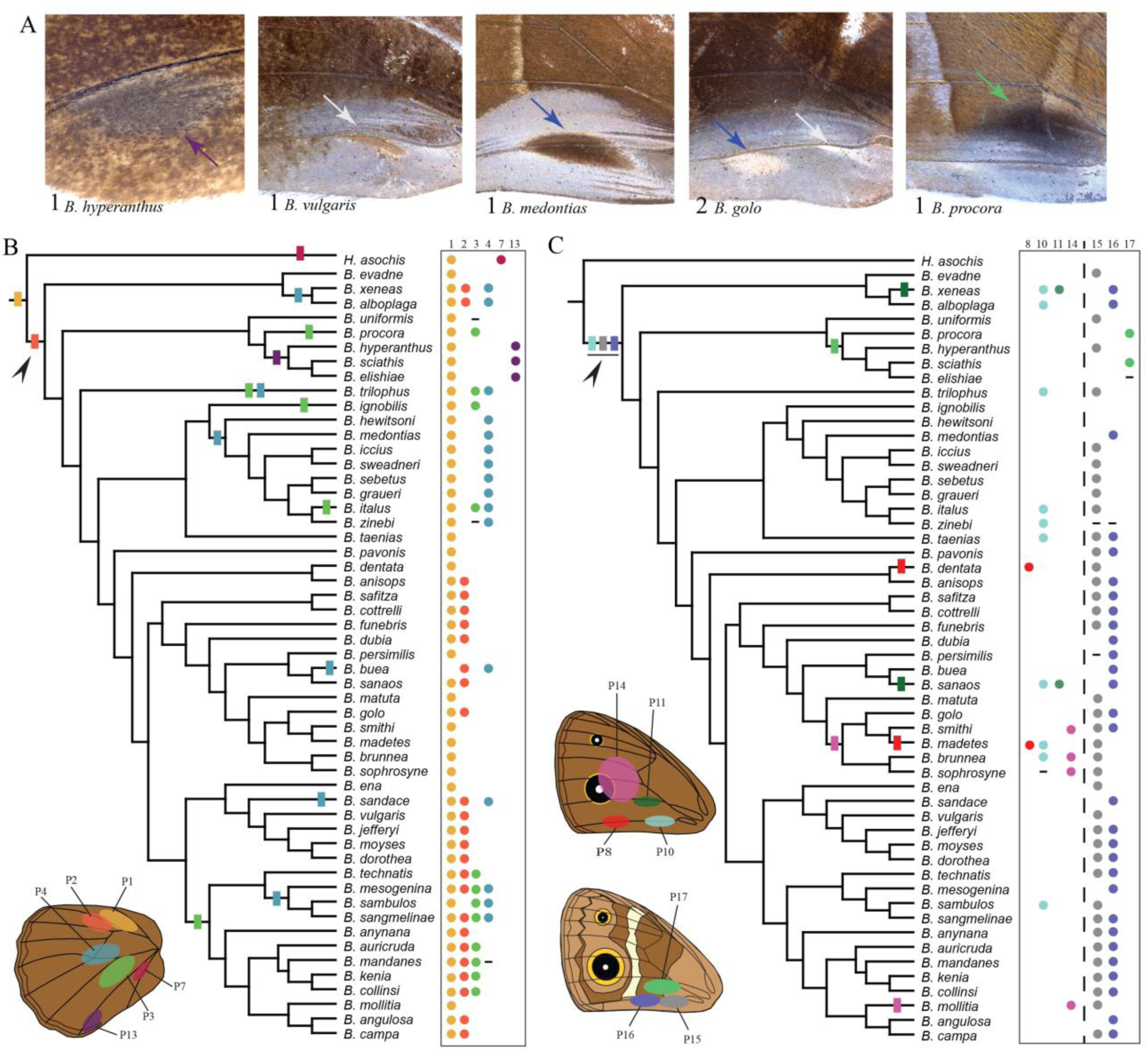
Diversity and evolution of patches on the wings of *Bicyclus* butterflies. A) From left: An example of patch 13 on the dorsal hindwing and combinations of patches 15, 16 and 17 on the ventral forewing of male *Bicyclus* butterflies. Note the broad area of silvery scales near the posterior margin of the ventral forewings. The number of patches and the name of the species is denoted at the bottom left. Colored arrows point to each patch and correspond to the colors used in the schematics in B and C. For examples of hindwing patches, refer to Figure 1B. B,C) Evolutionary history of (B) hindwing and (C) forewing scent patches at homologous positions on the wing within the *Bicyclus* lineage. Locations, number and color codes of the different patches are given in the schematic and their presence/absence in each species is listed to the right of the phylogenetic trees. Species for which data was not available are marked with ‘-’. Color codes of the patches match the codes of the corresponding hair-pencils in those sectors. Filled rectangles on the phylogenetic tree indicate the likely gains of the traits for which either a single or a multiple-origin scenario at the MRCA of all the species bearing that trait was significantly supported. For traits where a single- or multiple-origins scenario was equally supported, a single origin was mapped (black arrowheads).

Research into chemical ecology has spurred many studies on the chemical composition of pheromones, their plasticity and their evolution across the genus *Bicyclus* (Nymphalidae, Satyrinae, Satyrini) (Nieberding *et al.* 2008, 2012; Heuskin *et al.* 2014; Bacquet *et al.* 2015; Dion *et al.* 2016; Darragh *et al.* 2017; Balmer *et al.* 2018). However, no study to date has addressed the development and evolution of these sex-specific complexes. The male-specific androconia of *Bicyclus* butterflies are so diverse and variable in number, size, shape, color, chemical composition and position that they are a key trait in species identification (Condamin 1973; Brattström *et al.* 2015, 2016). The frequent changes in the morphology and location of the two components of the androconia – the hair-pencils and scent patches – within the *Bicyclus* clade, combined with its presence in the model species *Bicyclus anynana*, makes this a unique study system to understand the evolutionary history and, dissect the molecular and developmental genetic mechanisms governing male-specific pheromone complex development.

We used a combination of phylogenetics and a focused molecular investigation on the sex-determination gene, *doublesex (dsx)*, to address these questions. Our focus on *dsx* was due to two main reasons. Firstly, previous work on sex-specific trait development in *B. anynana* identified a non-cell-autonomous, hormonal mechanism as a determinant of sex-specific eyespot sizes (Bhardwaj *et al.* 2018). Here, sex-specific levels of the hormone 20-hydroxyecdysone and the presence of its receptor EcR cued the development of different dorsal eyespot sizes in males and females (Bhardwaj *et al.* 2018) but showed no effect on the development of the male-specific scent organs, indicating a different mechanism of determination of this sexually dimorphic trait. Secondly, in the rapidly increasing literature of sexually dimorphic trait development across insects, *dsx* appears to hold a highly conserved position in directing sex-specific gene expression that results in sex-specific morphologies and behaviors, cell-autonomously or in concert with hormones to incorporate environmental cues (Rideout *et al.* 2010; Kopp 2012; Gotoh *et al.* 2014; Prakash & Monteiro 2016). The *dsx* gene is spatially regulated in different somatic tissues such that only a subset of cells express *dsx* and are sex-aware (Robinett *et al.* 2010). It exhibits a modular genetic architecture, where modules evolve under different selective pressures, suggestive of functional partitioning between the different modules over evolutionary time (Baral *et al.* 2019). Moreover, distinct isoforms of *dsx* are produced in each sex, creating a sex and tissue-specific transcription factor asymmetry that can be used to direct cells into distinct developmental fates (Burtis & Baker 1989; Ito *et al.* 2013; Neville *et al.* 2014). Thus, evolution in patterns of spatial and temporal regulation of *dsx* (Tanaka *et al.* 2011), of its downstream targets (Ledón-Rettig *et al.* 2017), or in the mode of action of the different isoforms, i.e, as repressors or activators (Kopp *et al.* 2000; Kijimoto *et al.* 2012; Arbeitman *et al.* 2016), are all previously investigated mechanisms that can create vastly different, dynamically evolving, sex-specific morphologies in different insect lineages.

In this study, we first independently mapped the presence of hair-pencils and scent patches at homologous positions onto a phylogeny of the *Bicyclus* clade to understand the evolution of these traits. Then, we studied the role of *dsx* in the development of the sexually dimorphic scent organ complex in *Bicyclus anynana* using immunostainings and targeted gene knockouts using CRISPR/Cas9. Finally, we examined an outgroup species, *Orsotriaenae medus* (Nymphalidae, Satyrinae, Satyrini), that diverged from *Bicyclus* nearly 40 MYA, to estimate the degree of conservation of Dsx expression in scent organ development across satyrids.

## Results

### Androconia diversity within *Bicyclus*

Males in the genus *Bicyclus* display enormous diversity in the number, and position of the hair-pencils and scent patches on their wings (Fig 1B, Fig 2A). Hair-pencils are present only on the dorsal surfaces of both wings (Fig 1C, schematic) with the total number of forewing hair-pencils (0-2) being equal to or less than the number of hindwing hair-pencils (1-4) in any given species. Scent patches occur on all wing surfaces except the ventral hindwing (Fig 2B,C, schematic). As with the hair-pencils, forewing scent patches are rarer as compared to hindwing scent patches and vary from zero to two in number, per surface (dorsal/ventral) (Fig 1C, Fig 2C), while hindwings always have scent patches, varying in number from one to four (Fig 1C, Fig 2B). In most cases, hair-pencils are usually associated with an underlying patch (indicated by the same colors in Fig 1 and 2) but there are instances of hair-pencils present without any associated patch (hair-pencil 3 without patch 3 in *B. jefferyi, B. moyses* and *B. dorothea*) and vice versa (patch 13 in *B. hyperanthus, B. scaithis* and *B. elishiae*).

### Patterns of androconia diversity within *Bicyclus* occurs primarily via multiple trait gain, and trait loss

In order to understand the origins of this diversity within *Bicyclus*, we first constructed a phylogenetic tree of the species of interest using a Bayesian framework and then reconstructed the evolutionary history of the different hair-pencils and patches on the sampled trees. Our majority-rule Bayesian consensus tree (Supplementary Fig 1) was largely congruent with the Bayesian tree published in Aduse-Poku et al. 2017. We obtained the *Bicyclus* clade as a well-supported monophyletic group and the *evadne*-species group as a sister clade to all other *Bicyclus*, similar to the earlier published data. However, as with the Aduse-Poku et al. 2017 tree, some of the basal branches had poorer support giving rise to some basal uncertainty, which was accounted for in our ancestral state reconstructions.

We reconstructed the evolutionary history of the different hair-pencils and scent patches on the sampled trees. Ancestral state reconstruction of the hair-pencil present at the base of the discal cell, what we call hair-pencil 1, indicated that this trait is ancestral and was present before the divergence of the *Bicyclus* clade, with one loss in *Bicyclus buea* (Fig 1C yellow rectangle, Supplementary Table 2), with a similar situation for the associated patch 1 (Fig 2B yellow rectangle). Hypothesis tests for forewing hair-pencil 10 provided significant support for the model where the trait was present in the most recent common ancestor (MRCA), similar to hair-pencil and patch 1 (Fig 1C light blue rectangle, Supplementary Table 2) followed by multiple losses. For hair-pencils 2, 4 and 6 (Fig 1C orange, blue and lilac) and the patches 3 and 4 (Fig 2B green and blue), hypothesis tests comparing alternative reconstructions of trait origin within *Bicyclus* i.e. the definitive presence vs the definitive absence of the trait at the MRCA of all the species bearing the trait, provided significant support towards the model where the trait was absent in the MRCA (Supplementary Table 2). Significant support for MRCA=0 was also obtained for dorsal forewing patches 8, 11 and 14 (Fig 2C red, dark green and pink). This supports a scenario of multiple, independent gains of the different traits during the evolution of *Bicyclus* with subsequent losses in many lineages. To broadly understand the likely number of independent gains of these traits, we used stochastic character mapping and an unconstrained model of trait evolution to identify gains at internal nodes (Fig 1C and Fig 2B, C; rectangles). For example, hair-pencil 2 was likely gained at least twice – once in the *evadne*-species group and once after the branching off of the *ignobilis-hewitsoni* group (Fig 1C orange rectangles and arrows). Similarly, hair-pencil 4 was likely gained six times within *Bicyclus* (Fig 1C, blue rectangles and stars). However, for a few traits including hindwing hair-pencil 3 (Fig 1C), hindwing patch 2 (Fig 2B), dorsal forewing patch 10 (Fig 2C) and ventral forewing patches 15 and 16 (Fig 2C) there was no definitive support for one state over the other at the MRCA, making the reconstruction of the evolution of these traits ambiguous (Supplementary Table 2). For such traits, we adopted the most parsimonious evolutionary explanation of a single origin at the MRCA (Fig 1C, Fig 2B, C; black arrowheads). Trait losses are not mapped on the phylogenetic trees.

### Scent patches and hair-pencils appear to evolve in a correlated manner

We next tested for correlated evolution between pairs of hair-pencils and patches. Hair-pencils 1, 2, 3 and 4 and patches 1, 2, 3 and 4, respectively, showed strong support for dependent evolution (Supplementary Table 3) consistent with our understanding of a hair-pencil and a patch together making a functional composite scent organ for pheromone dissemination. In addition, there was positive support for correlated evolution between hair-pencil 1 and patch 15 and, very strong support for correlated evolution between hair-pencil 2 and patch 16 (Supplementary Table 3). Patches 15 and 16 which are located on the ventral forewing are in contact with hair-pencils 1 and 2 on the dorsal hindwing respectively at the region of overlap between the two wings. The evidence for correlated evolution between these forewing patches and hindwing hair-pencils are again suggestive of a composite scent organ, now dispersed between two different surfaces.

### Dsx expression changes from monomorphic to sex-specific in the presumptive scent organ wing regions over the course of development

To identify the proximate mechanisms for male-specific androconia development, we examined the expression of the sex-determinant protein Dsx in developing wing discs of both sexes in *B. anynana*. Adult male *B. anynana* butterflies possess dorsal hindwing hair-pencils 1 and 2 with their corresponding patches, and ventral forewing patches 15 and 16. Secretory cells lie beneath dorsal hindwing patch 1 and ventral forewing patches 15 and 16 (Dion *et al.* 2016). In addition, males also possess the broad area of silvery scales near the posterior margin of the ventral forewing. Patterns of Dsx expression mapped to the general scent patch and hair-pencil regions early in wing development and were initially similar on the forewing and hindwing discs of both sexes (Fig. 3, Mid-5th instar). As development progressed, Dsx showed sex-specific variation in the presumptive scent organ regions: from the last larval instar (5^th^), through the wandering stage, to the pupal stage (Fig 3). The pre-pupal stage was not conducive to any sort of staining, and inferences about the role of Dsx at this stage were based on Dsx expression in the preceding and succeeding developmental stages and will be discussed later. Forewing and hindwing Dsx expression patterns are described below.

**Figure 3:**
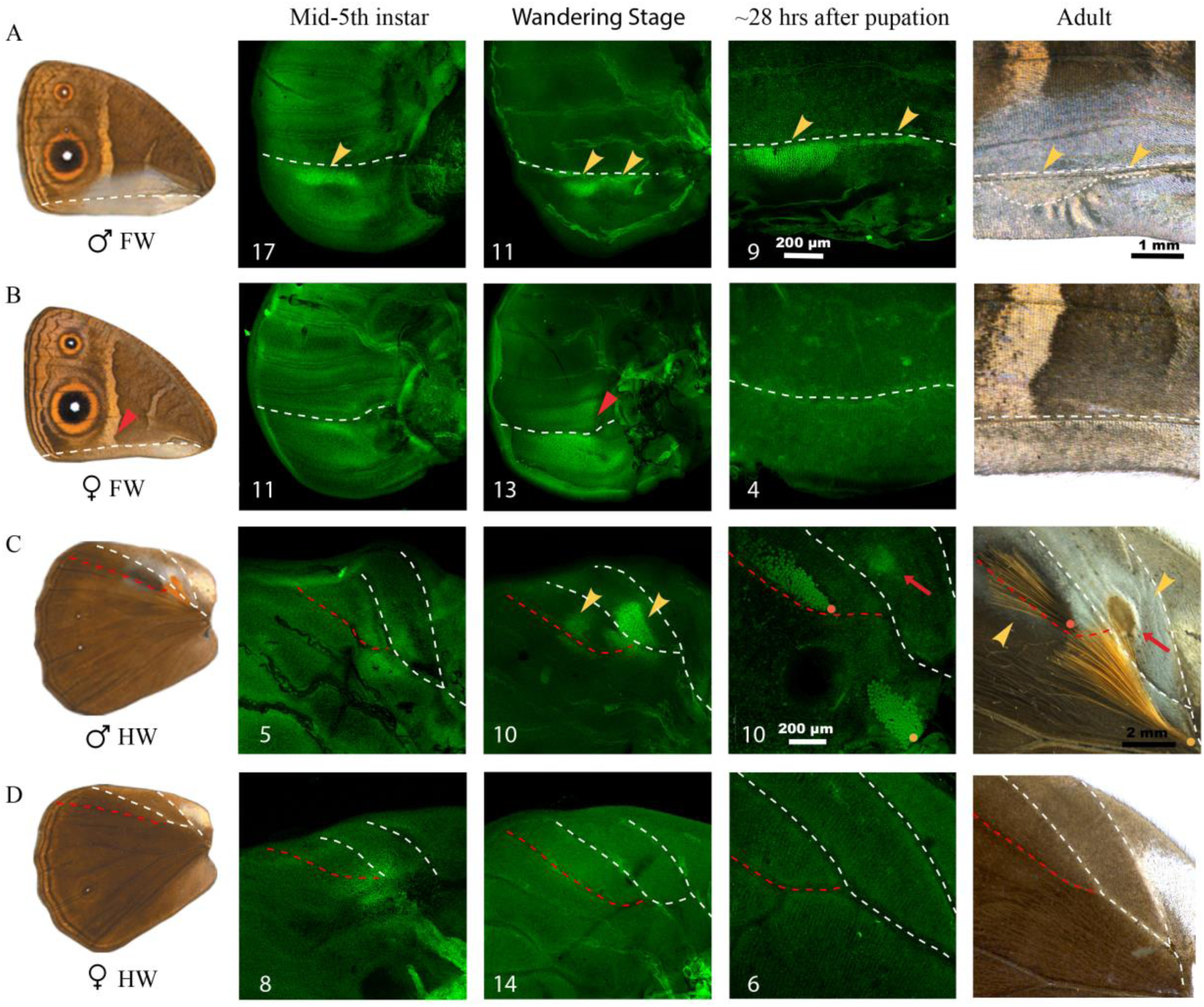
Dsx expression during wing development in *Bicyclus anynana*. Dsx expression during three different wing developmental stages i.e. mid-5^th^ instar, wandering stage and 28 hours after pupation are shown for (A) male and (B) female forewings and (C) male and (D) female hindwings. The adult wings are shown in the first column and an expanded view of the scent organ regions in both sexes is shown in the last column. Images shown are the best, illustrative images and numbers at the bottom left indicate sample size. White and red dotted lines denote homologous veins in the respective wings and the orange and yellow dots in (C) correspond to the base of hair-pencils as in Figure 1. Yellow arrowheads in (A) and (C) indicate the two scent patches on the forewing and hindwing respectively and the red arrows in (C) indicate the secretory cells. Scale bars are shown for the pupal wings and adult structures.

The Dsx expression pattern in forewings became associated with the development of the scent patches and silver scales in males. In mid-5^th^ instar forewing discs, Dsx was expressed in sector 1A+2A in both sexes (sector and vein nomenclature in Supplementary Fig 2), with a stronger, narrow linear expression in the anterior part of the sector (Fig 3A,B; below the white dotted line - yellow arrowhead). In the following wandering stage, expression diverged across the sexes. In males, the single, linear expression of Dsx was defined into two domains, corresponding to the future locations of scent patches 15 and 16, which was further refined into precise spatial domains, resembling the shapes of the two patches, about 28 hours after pupation (Fig 3A; yellow arrowheads). At this pupal stage Dsx expression was also present in punctate nuclei that mapped to the general region of the ventral forewing silver scales in males. In females, however, expression of Dsx remained broad within the proximal sector 1A+2A, expanded into the region of the white band in the Cu2 sector (Figure 3B; red arrowhead), but, by 28 hours after pupation, no Dsx expression was seen in the corresponding wing regions (Fig 3B).

The hindwing discs also showed a similar progression of Dsx expression over time where the initial broad association of Dsx with the location of both scent patches and hair-pencils in both sexes became more refined and restricted in male wings only. In both male and female mid-5^th^ instar discs, there was a circular region of Dsx expression at the proximal end of sector Rs, extending partially into the discal cell (Fig 3C,D), which potentially covered both the scent patch and hair-pencil domains. This broad expression domain was later refined into two spatial regions in male wandering stage discs corresponding to the male hindwing scent patches with surrounding silver scales (patch 1) and, the patch of greyish-silver scales that lie below the future black hair-pencils (patch 2) (Fig 3C; yellow arrowheads). No Dsx expression was seen in the presumptive hair-pencils at this stage. About a day after pupation, male hindwing expression of Dsx was spatially refined and modified, now marking the secretory cells underneath patch 1 (Fig 3C; red arrows) and both hair-pencils (Fig 3C). As mentioned previously, secretory cells underly certain patches in *Bicyclus* species, including *B. anynana*, but not all patches. In females, the initial expression of Dsx was lost in the wandering stage and by 28 hours after pupation there was no Dsx expression in corresponding regions of the female hindwing discs (Fig 3D).

### Sex-specific isoforms of *dsx* regulate the development of the two components of the scent organ in different ways

To identify if *dsx* was sex-specifically spliced in the male and female wing tissues of *B. anynana, dsx* was amplified and sequenced from the wings. Male and female pupal wing discs expressed different isoforms of *dsx* (Supplementary Fig 3) suggesting that *dsx* is indeed sex-specifically spliced in the wing tissues. The partial sequences generated are available on GenBank (accession nos. MK869725 and MK869726).

To functionally verify the role of *dsx* in the development of the male-specific scent organs in *B. anynana*, we used CRISPR/Cas9, targeting the common DNA binding domain, which is shared between the male- and female-specific isoforms (Fig 4A). In males, knockout of *dsxM* affected the development of both forewing and hindwing scent patches and surrounding silver scales, corresponding to Dsx’s region of expression in developing wing discs. For both wings, crispants showed varying degrees of loss of silver scales (Fig 4B, C; red arrowheads) and scent patches with secretory cells (Fig 4C, D, E; red arrowheads, Supplementary Fig 4B, C; black arrows). These results suggest an activating role for *dsxM* in males, in the development of the scent patches and the broad silvery scale region. Most interestingly, however, despite the loss of the greyish-silver scales (patch 2) that lie beneath the black hair-pencil (Supplementary Fig 4D; black arrowheads), indicating effective disruption of *dsx* in this wing area, there was never a loss of either the black or yellow hair-pencils, though hair-pencil morphology, color and density were affected in many individuals (Supplementary Fig 4D).

**Figure 4:**
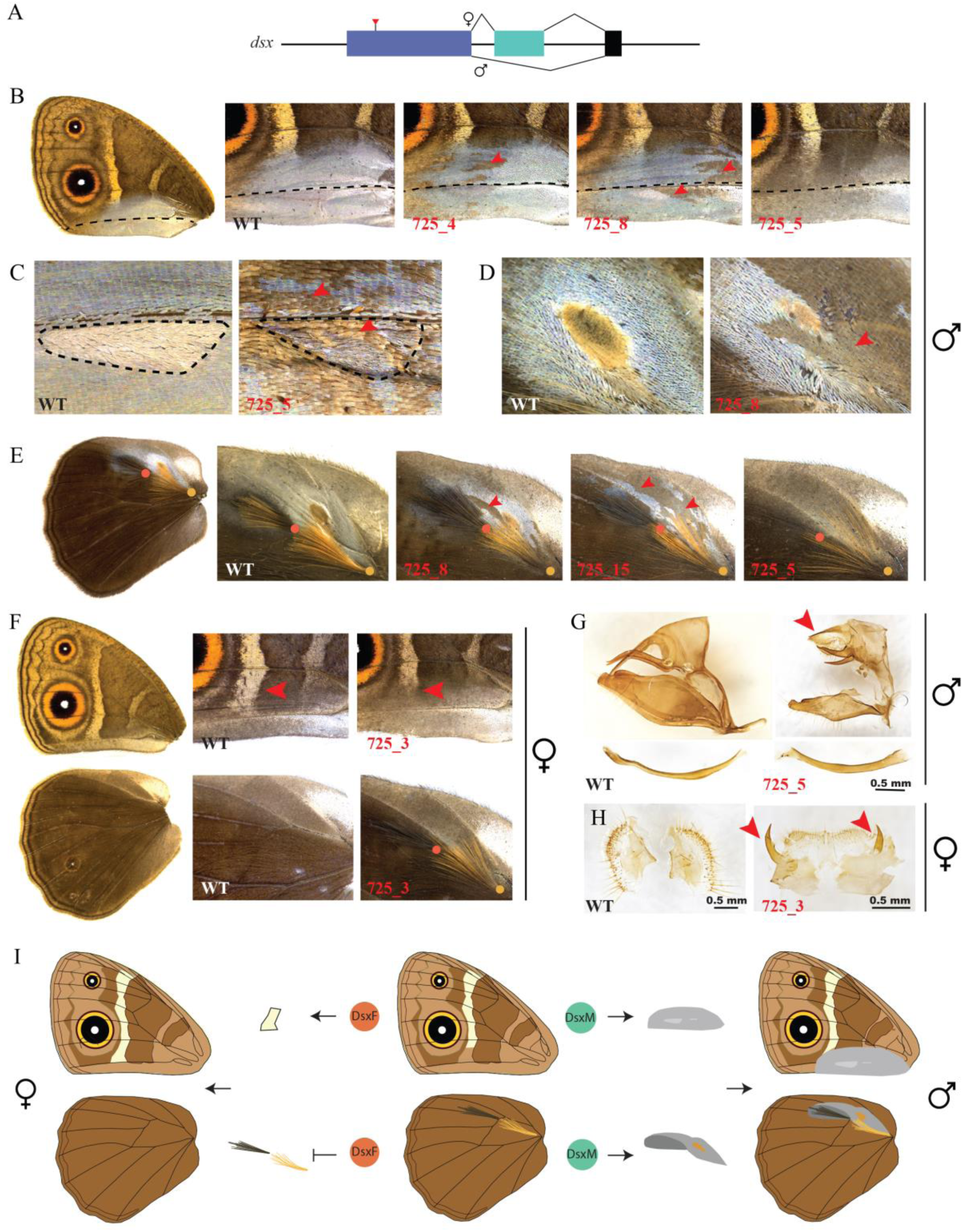
CRISPR/Cas9 directed *dsx* crispant phenotypes in *Bicyclus anynana*. A) Schematic of *dsx* male and female isoforms with the region targeted by CRISPR/Cas9 marked with a red arrow. In panels B-H, wildtype (WT) is shown to the left and crispants are indicated in red to the right. (B-E) Mosaic phenotypes on male (B,C) ventral forewing patches and broad silvery scale region and (D,E) dorsal hindwing patches. Black dotted lines in (B) highlight homologous veins and in (C) outlines patch 16. Red arrowheads indicate some of the mosaic phenotypes F) *dsx* crispant phenotypes on female ventral forewings (top) and dorsal hindwings (bottom). Red arrowheads indicate the differences in length of the white band between WT and crispant. G) Intersex phenotypes of male genitalia with reduced claspers, aedeagus (bottom) and an intermediate morphology of the uncus (red arrowhead) H) Intersex phenotypes of female genitalia. Intermediate structures are indicated with red arrowheads I) Schematic of the effects of *dsx* on *B. anynana* wing patterns. Centre shows the wing phenotype of ventral forewing (top) and dorsal hindwing (bottom) in the absence of either *dsx* isoform. Dsx in males activates the development of patches on both fore and hindwings (right) while Dsx in females increases the length of the white band towards the posterior end of the forewing and represses hair-pencils on the hindwing (left).

Knockout of *dsxF* in females showed contrasting results on its effect on the two androconia components. Crispant females expressed pairs of dorsal hair-pencils on either both or only one hindwing. These hair-pencils were mostly normal but had a lower density of hair-like scales (Fig 4F, Supplementary Fig 4E), suggesting that *dsx* in females represses hair-pencil development (Fig 4I). However, none of the female crispants displayed any patches and associated silver scales on either forewings or hindwings, indicating the lack of a role of *dsxF* in repressing the development of these scent organ components (Fig 4I). Additionally, on the ventral forewings, *dsx* crispant females showed modifications to the white transversal band that now stopped midway into the CuA2 sector instead of extending all the way to vein 1A+2A as in wildtype (Fig 4F; red arrowheads). This indicates that *dsxF* promotes the development of the posterior band in females. None of the *dsx* crispants showed any effects on eyespot size. Injection statistics and quantification of the different *dsx* crispant phenotypes is provided in Supplementary Tables 5 and 6.

### *dsx* affects the morphology of the genitalia

*dsx* crispant males and females also developed deformed genitalia, displaying intersex phenotypes. The aedeagus and claspers in males became reduced in size, while the main body of the genital structure, the uncus, displayed an intermediate morphology (Fig 4G; red arrowhead). In females, the two sclerotized genital plates also developed intersex structures (Fig 4H). The sex of crispants was verified by amplifying a W-microsatellite that only occurs in females (Supplementary Fig 4F) and effective *dsx* disruptions were confirmed by sequencing DNA from thoracic tissue (Supplementary Fig 4A).

### Scent organ associated Dsx expression arose before the divergence of the *Bicyclus* lineage

To verify if *dsx*-mediated development of androconia in *B. anynana* and, potentially all other *Bicyclus* species predated the origin of the genus, we performed Dsx antibody stainings on mid-5^th^ instar forewing discs of a distantly related satyrid species, *Orsotriaena medus*, displaying male-specific scent organs. Males of this species possess a pocket-like scent patch on the dorsal forewings with an associated hair-pencil, that appears as an extruded portion of the wing on the ventral forewing surface (Fig 5A Adult; yellow arrowhead indicating the extruded wing patch on the ventral forewing). Both male and female mid-5^th^ instar forewing discs showed spatial expression of Dsx in the presumptive scent organ regions (Fig 5). This could map to either hair-pencil or patch in males but is unclear at this time because we did not look at surface-specific expression of Dsx i.e. dorsal or ventral expression. Nevertheless, these observations, similar to those in *B. anyanna*, suggest that scent organ associated Dsx expression likely originated before the divergence of the *Bicyclus* and *Orsotriaena* lineages.

**Figure 5:**
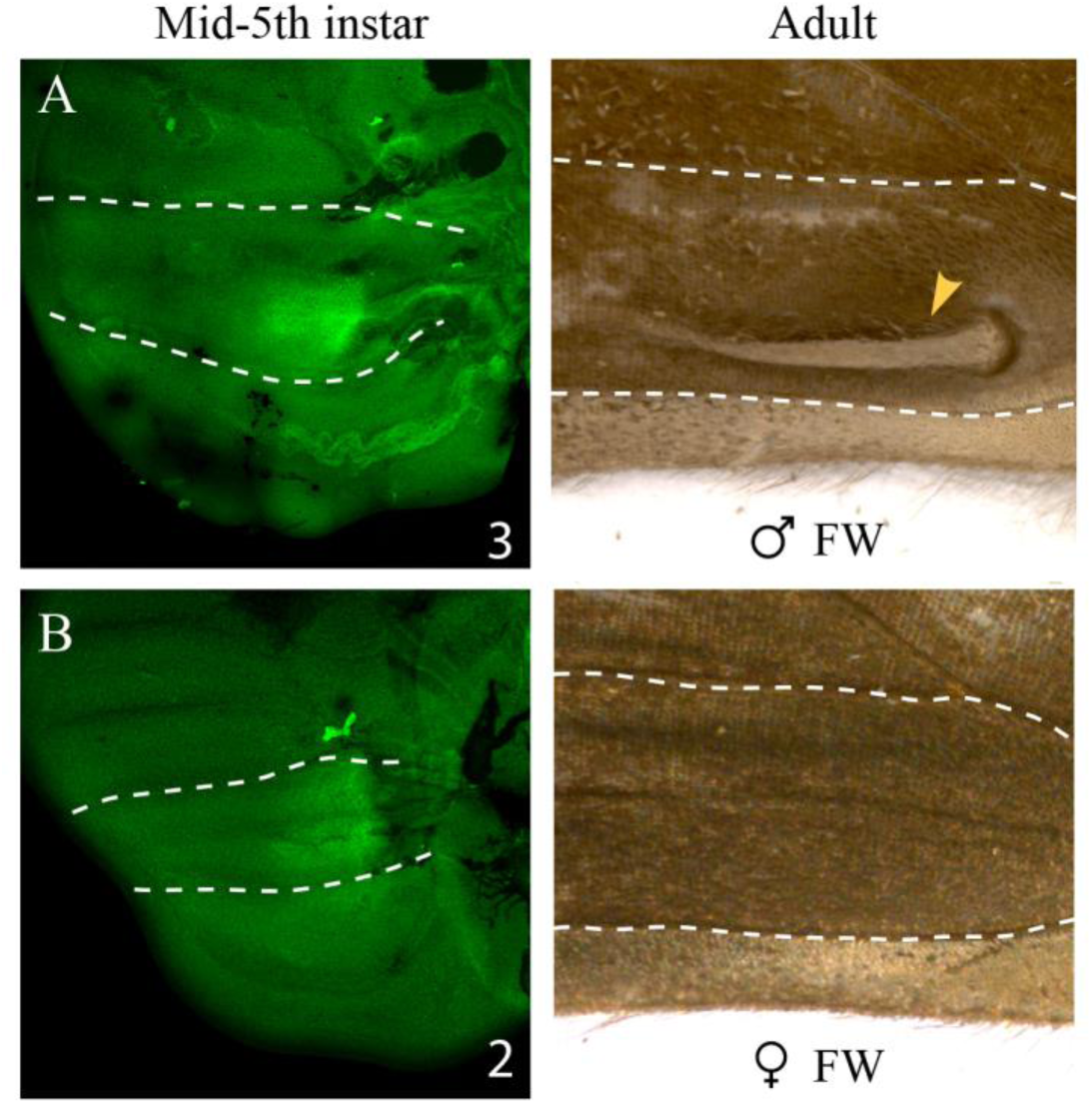
Dsx expression in developing forewing discs of *Orsotriaena medus*. Dsx expression (left) in mid-5^th^ instar forewing discs of (A) males and (B) females. Corresponding adult ventral wing regions are shown to the right. Note the extruded scent patch present only in adult male wings (yellow arrowhead). Images shown are the best, illustrative images and numbers at the bottom right indicate sample size. White lines indicate homologous veins in the respective wings.

## Discussion

### Patterns of androconia evolution within *Bicyclus*

Our phylogenetic analyses suggest that androconia have extremely labile evolutionary histories, with frequent gains and losses of both scent patches and hair-pencils - a pattern of evolution not unlike that of dorsal eyespots, which are also extremely labile within *Bicyclus* (Oliver *et al.* 2009). Most of the traits investigated were absent in the common ancestor to all *Bicyclus*, except for the hair-pencil 1 at the base of the discal cell on the hindwings and its corresponding patch 1, which are both old and evolutionarily stable. Most other hair-pencils and patches were gained multiple times during the evolution of this genus. This pattern suggests that the gene regulatory networks (GRNs) that build hair-pencils and patches were kept intact throughout the *Bicyclus* radiation, because one or more of these traits were always present somewhere on the wings, but redeployments and losses of the GRNs took place during evolution. Evolution in trait number might perhaps be achieved via tinkering with the expression of master regulator genes of either GRN, still to be discovered. Thus the evolution of the composite scent organs in *Bicyclus* provide additional evidence against Dollo’s law (the idea that complex traits once lost in evolution cannot be regained) and show evolutionary trends similar to loss and re-gain of other complex characters such as eyespots (Oliver *et al.* 2009) or shell coiling in gastropods (Collin & Cipriani 2003).

Additionally, hair-pencils and patches also showed correlated evolution across *Bicyclus*. This makes sense considering that the two components together form a functional composite organ required for pheromone production and dissemination in some species such as *B. anynana*. However, many species possess only one of these functionally coupled traits which begs the question of their functional roles in these species. Patches without correspondent hair-pencils were more commonly present than hair-pencils without correspondent patches. Examples for both include the presence of velvety patches on the dorsal forewings and hindwings of members of the *martius*- and *sciathis*-groups respectively (Fig 2, patches 13 and 14), or the presence of hair-pencil 3 and the absence of corresponding patch 3 in *B. jefferyi, B. moyses* and *B. dorothea* (Fig 1C and Fig 2B). Despite the possibility of patches being functional in chemical communication, independently of associated hair-pencils, we note that many patches in *Bicyclus* do not secrete any pheromones (Bacquet *et al.* 2015). Thus, the functionality of these organs and their role, or lack of it, in the behavioral biology of *Bicyclus* butterflies still remains an open question. Their functions in chemical communication might have been lost over evolutionary time or these organs might now function in visual communication instead (Bacquet *et al.* 2015).

Since our analyses were limited to the *Bicyclus* clade, it is still unclear how many other lineages possess scent organs, and where (which lineage and which position on the wing) the first hair-pencil and patch serial homolog originated. A more detailed and inclusive phylogenetic sampling across all lineages with scent organs would be needed to determine the origins of these traits and compare their evolutionary patterns with what we identified in the *Bicyclus* clade.

### Precise and refined spatial expression of Dsx determines sex-specific scent organs

Investigation of the proximate mechanisms governing male-specific scent organ development in *B. anynana*, identified two different modes of action of the sex-specific *dsx* isoforms in determining the two components of the organ. Precise spatial expression of Dsx on male wings preceded the development of male-specific scent patches. This expression domain, initially broad and non-sex-specific, was refined during development to map precisely to the final adult male structures. Dsx expression was also visualized in the silvery scale region on the ventral forewings at early stages of pupation, and our functional data indicates that this gene is also required for the development of these scales. In contrast, the development of hair-pencils in males seems to be largely Dsx independent, as hair-pencils can develop in both sexes when *dsx* is disrupted. Instead, Dsx in females appears to be functioning as a repressing factor that prevents hair-pencils from forming in females. The expression of this gene in the general area of the hair-pencils early in wing development, rather than in the precise location of the hair pencils in female wings, could be sufficient for its repressive function. On the other hand, spatially refined Dsx expression patterns in female wings may have been present in the pre-pupal stage, which was not investigated in this study. These results suggest that the two components of the scent organ in *B. anynana* are determined at different developmental stages via modulation of an initially non-sex-specific expression of Dsx, and that the sex-limited presence of this organ is governed by two different modes of *dsx* action: *dsxM* acting as an activator of scent patches in males and *dsxF* acting as a repressor of hair-pencils in females, modes that was previously identified in *Drosophila melanogaster* (Goldman & Arbeitman 2007; Arbeitman *et al.* 2016). Such distinct modes of action of *dsx* isoforms have also been identified in other species. For example, in the beetle *Onthophagus taurus*, the male isoform of *doublesex* promotes head horn development while the female isoform represses horn development in females (Kijimoto *et al.* 2012). In contrast, in a closely related species, *Onthophagus sagittarius*, that exhibits a reversed sex-specific thoracic horn phenotype, the male *doublesex* isoform inhibits horn development while the female isoform promotes its growth (Kijimoto *et al.* 2012). Furthermore, in the case of *Drosophila* abdominal pigmentation, while the male isoform has a negligible effect on pigmentation, the female isoform represses such pigmentation in females (Kopp *et al.* 2000).

The sex-specific development of the scent organs is determined by the spatial regulation of *dsx*, in a similar way to the regulation of *dsx* in the sex-combs in *Drosophila* (Tanaka *et al.* 2011). Spatial regulation of *dsxM* is required for the precise development of the male-specific patches while potentially less precise spatial regulation of *dsxF* is required for the repression of the hair-pencils in females, leading to a male-limited occurrence of this trait. The refinement of Dsx expression in males suggests the involvement of other spatially-restricted transcription factors in regulating *dsx* over time, similar to the *Scr*-*dsx* feedback loop involved in *Drosophila* sex-comb development (Tanaka *et al.* 2011). Candidate transcription factors that might be involved in dorsal surface-specific expression of *dsx* include *apterous A* and factors downstream of *apterous A* (Prakash & Monteiro 2018), whereas ventral-specific factors are unknown. In addition, the downstream targets of *dsx* in the development of the scent organs still remains unanswered and is an interesting avenue for future research.

Further, our identification of scent organ associated Dsx expression in the more distantly related satyrid butterfly *Orsotriaena medus* suggests that *dsx*-mediated sex-specific androconial development arose before the divergence of these two clades. It is then highly plausible that the proximate mechanisms we have identified here in *B. anynana* are applicable to other *Bicyclus* species, with the male-specific *dsx* isoform required only for patch development in males while the female *dsx* isoform is required to repress hair-pencils in females. This hypothesis, however, requires additional functional work in the other *Bicyclus* species.

### Molecular mechanisms of scent organ development and the evolution of androconia diversity

Given our understanding of the proximate mechanisms of sex-specific scent organ development in one *Bicyclus* species, and the patterns of trait evolution within this genus, we can draw certain inferences and propose hypotheses about the mechanisms of androconia evolution within the *Bicyclus* clade. Since spatial regulation of the male *dsx* isoform is required for patch development, diversification in the number of patches could have occurred via the gain and loss of new domains of *dsx* expression on the wing. However, the diversity in hair-pencils cannot be explained via the acquisition and loss of new domains of *dsx* because hair-pencil development can occur independently of *dsx*. Instead, we hypothesize that the origin of the hair-pencil GRN probably arose via the deployment of the GRN that determines long hair-like scales, in a dense cluster of neighboring cells. These long scales are morphologically like hair-pencils and occur at low density on the wings of many butterflies of both sexes, even in families that do not possess hair-pencils (Supplementary Fig 5). The hair-pencil GRN could then have been modified and co-opted to different locations on the wing, generating diversity in number and morphology with sex-specificity being created by Dsx-mediated repression of this network in females. Thus, in the particular case of hair-pencils, spatial regulation of *dsx* does not lead to diversity in sex-specific traits, a situation that differs from that of sex-comb evolution in *Drosophila* (Tanaka *et al.* 2011).

An additional possibility for the diversity in scent organs observed could be due to introgression or the process of gene transfer between closely related species because of hybridization. Wing pattern mimicry between closely related *Heliconius* species has previously been explained by adaptive introgression (The Heliconius Genome Consortium 2012; Smith & Kronforst 2013; Zhang *et al.* 2016) and such a mechanism could also explain diversity in androconia within *Bicyclus*, where many species occur in sympatry in Africa. Identifying the on and off switches of key genes in the GRNs that create hair-pencils and patches and, performing comparative molecular studies can help validate these hypotheses and also potentially explain the correlated evolution between the two traits.

### Different proximate mechanisms generate male-specific structures on the same wing surface

Our results show two different mechanisms governing the development of sex-specific traits within one species, bearing resemblance to both Darwin and Wallace’s views on the origin of sexual trait dimorphism. *dsxM* leads to activation of scent patches in males only, lending support to Darwin’s theory of a trait arising in a single sex right from the very beginning. On the other hand, *dsxF* represses hair-pencil development in females, suggesting that the hair-pencil GRN might have initially originated in both sexes, but was subsequently removed from females, resembling Wallace’s origins of sexually dimorphic traits. However, we acknowledge that both the Darwin and Wallace views on the origins of sexual trait dimorphisms, in this particular case of scent organ origins, would need further testing by functional verification in other *Bicyclus* species.

This study, concurrent with the study on sex-specific eyespot development in *B. anynana* (Bhardwaj *et al.* 2018), have together identified three different ways of producing sexual dimorphisms in one species via both cell-autonomous and non-cell-autonomous mechanisms – activation of the scent patches only in males via DsxM, female-specific repression of hair-pencils via DsxF (both acting through cell-autonomous mechanisms) and, a third, non-autonomous hormonal threshold mechanism controlling sex-specific eyespot sizes. This diversity in proximate mechanisms within one organism has implications for the origins and rapid turnover of sexually dimorphic traits and must be considered while understanding the evolution of sex-specific traits.

## Materials and methods

### Animal husbandry

Mated *Orsotriaena medus* females were collected in forested areas of Singapore under the permit number NP/RP14-063-3a. *Bicyclus anynana* and *Orsotriaena medus* butterflies were reared in a 27°C temperature-controlled room with 65% humidity and a 12:12 hour light:dark photoperiod. Adults of both species were fed on banana. *B. anynana* larvae were fed on young corn plants while *O. medus* larvae were fed on *Ischaemum sp.* grasses, commonly found in Singapore.

### Character sampling and phylogenetic analyses

53 *Bicyclus* species listed in the phylogeny of Monteiro and Pierce (Monteiro & Pierce 2001) and 1 outgroup were considered for this study, however, we used the larger set of 179 individuals (95 *Bicyclus* species and 10 outgroups), 10 gene dataset from Aduse-Poku et al. 2017 to reconstruct a Bayesian phylogenetic tree using MrBayes v3.2.7a (Ronquist *et al.* 2012). This reconstruction allowed us to account for phylogenetic uncertainty in the ancestral state reconstruction of the androconia. Nucleotide sequences were aligned based on the translated amino acid sequences and then concatenated to create a dataset of 7704 characters from 10 genes. The partitioning scheme provided by Aduse-Poku et al 2017 was used to partition this dataset. Two parallels of four chains (three heated and one cold) was run on MrBayes for 5 million generations and a split frequency below 0.01 was used to assess stationarity. Burn-in was set to 25% (2500 trees) and the remaining set of trees from both runs were pooled together to total 15000 trees for the following analyses. The postburn-in trees were also used to produce a majority-rule consensus tree. These trees were then pruned in Mesquite (Maddison & Maddison 2018) to only include the 54 taxa used in the Monteiro and Pierce study, which we had in hand and could examine for the presence and location of the male scent organs.

For character sampling, hindwings and forewings of the different *Bicyclus* and outgroup species were imaged using a Leica DMS 1000 microscope. The images in combination with data from Condamin (Condamin 1973) and other scientific literature (Nieberding *et al.* 2008; Bacquet *et al.* 2015; Brattström *et al.* 2015, 2016; Aduse-Poku *et al.* 2017) were used to score the presence and absence of hair-pencils and scent patches in each species. We defined a hair-pencil as a group of tightly packed brush-like scales. On the dorsal hindwing and dorsal forewing, we defined a scent patch as an area of modified epidermal scales, different from the background color. We did not examine the presence or absence of secretory cells beneath the patches. On the ventral forewing, *Bicyclus* species usually possess a broad area of silvery scales near the posterior margin of the wing. Within this region there are usually one to two smaller areas of modified epidermal scales, most often associated with secretory cells underneath. We scored these smaller regions as scent patches on the ventral forewing, similar to the scoring scheme in Bacquet et al. 2015. In cases where two scent patches were located within sector 1A+2A (see sector and vein nomenclature in Supplementary Fig 2) on the ventral forewing, they were distinguished based on their position to the left or right side of a hypothetical perpendicular line drawn from the intersection of veins M3 and CuA1 to the vein 1A+2A. Hair-pencils and patches were considered homologous between species if they occupied similar positions within the same sector on the wing, i.e, a region bound by the wing veins. The character matrix is provided in Supplementary Table 1.

### Ancestral state reconstruction

Visualizations of the ancestral state reconstructions of the different traits was carried out in Mesquite (Maddison & Maddison 2018) using the majority-rule consensus tree. A two-parameter asymmetric model of evolution was used to allow for different rates of gain and loss of hair-pencils and patches.

For a more rigorous statistical hypothesis testing of the origins of the different traits within *Bicyclus*, ancestral states were also reconstructed across the posterior distribution of trees generated in MrBayes (15000 trees) using a reverse jump MCMC method as implemented in the MULTISTATE package in BayesTraits V3 (http://www.evolution.rdg.ac.uk/BayesTraitsV3.0.1/BayesTraitsV3.0.1.html) (Pagel *et al.* 2004). Reverse jump MCMC accounts for both phylogenetic uncertainty and the uncertainty in estimation of parameters of the model for trait evolution during the ancestral state reconstruction. To test whether a particular state i.e. 1 (trait present) or 0 (trait absent), was statistically supported for each trait at the most recent common ancestor (MRCA) of all lineages bearing the trait of interest, model marginal likelihoods were calculated with each alternative state fixed at the MRCA. Phylogenetic hypothesis testing was done by comparing the log marginal likelihoods of the two models and a model was considered significantly more likely if the log Bayes Factor = 2*Δ log marginal likelihood, was greater than 2 (Pagel 1999). When the trait was present at the MRCA of the species bearing it, then it was considered homologous, otherwise, it was considered analogous (with multiple origins). All RJ-MCMC chains were run for 5 million generations with a burn-in of 25% and a uniform prior between 0 and 100. The marginal likelihoods of the different models were estimated using 500 stones of a stepping stone sampler as per the BayesTraits manual. Further, to map the gains of traits that were significantly supported as either homologous or not within *Bicyclus*, we ran an unconstrainted model of trait evolution and estimated the probable ancestral states at internal nodes. We also ran stochastic character mapping in Mesquite and used both the analyses to broadly understand the evolution of hair-pencils and patches in *Bicyclus*.

Correlations between pairs of traits was estimated using the program DISCRETE in BayesTraits (Pagel & Meade 2006), with a RJ-MCMC run across the 15000 postburn-in trees and the same parameters as mentioned above. Dependence between two traits were investigated by comparing the log marginal likelihoods of an independent model (traits evolve independently) vs a dependent model (traits evolve in a correlated manner) and a log Bayes Factor = 2*(log marginal likelihood (dependent model) – log marginal likelihood (independent model)) >2 was considered a significant support towards the dependent model.

### Dsx immunostainings

We used a primary monoclonal antibody (mouse) raised against the *Drosophila* Dsx protein DNA binding domain (DsxDBD; Developmental Studies Hybridoma Bank deposition by Baker, B.S) (Mellert *et al.* 2012) that is present in both male and female isoforms of Dsx. This antibody was previously used to detect sex-specific Dsx expression on *Agraulis vanilla* (Martin *et al.* 2014) and *Papilio polytes* wing discs (Kunte *et al.* 2014). Alexa Fluor 488-conjugated donkey anti-mouse antibody (Jackson ImmunoResearch Laboratories, Inc) was used as a secondary antibody.

Wing discs were dissected from both sexes at different stages during development and transferred to cold fix buffer (0.1 M PIPES pH 6.9, 1 mM EGTA pH 6.9, 1% Triton x-100, 2 mM MgSO_4_). Pupal wings were transferred to fix buffer at room temperature and moved onto ice after addition of fixative to prevent crumpling of the tissue. Fixation was in 4% formaldehyde (added directly to the wells) for 30 min on ice, followed by 5 washes with PBS. For larval wings, the peripodial membrane was removed after fixation. The wings were then transferred to block buffer (50 mM Tris pH 6.8, 150 mM NaCl, 0.5% IGEPAL, 5 mg/ml BSA) overnight. Incubation with primary antibody at a concentration of 2.5 µg/ml in wash buffer (50 mM Tris pH 6.8, 150 mM NaCl, 0.5% IGEPAL, 1 mg/ml BSA) was for 1 hour at room temperature, followed by 4 washes with wash buffer. Wings were subsequently incubated in secondary antibody (1:500) diluted in wash buffer for 30 min at room temperature and then washed with wash buffer 8-10 times, with the final wash going overnight. Samples were mounted on slides with mounting media and imaged on an Olympus FLUOVIEW FV3000 confocal microscope.

### Identifying *doublesex* isoforms in wings

Total RNA was extracted from ∼18 hr pupal wings of males and females using TRIzol reagent (Invitrogen) according to the manufacturer’s protocol. cDNA was prepared by reverse transcription using the RevertAid First Strand cDNA Synthesis Kit (Thermo Scientific, USA). The canonical male and female *dsx* isoform sequences in *Bicyclus anynana* were kindly provided by Arjen Van’t Hof. *dsx* transcritps were amplified from the wing tissues of both sexes using primers designed against sequences that are common to both male and female isoforms (Supplementary Table 4). PCR amplifications were done using 2x PCRBIO Taq Red Mix (PCR Biosystems) with the following conditions: 1 min at 96°C, 40 cycles of 15s at 96°C, 15s at 60°C and 10s at 72°C.

### CRISPR/Cas9 gene editing

Cas9-mediated gene editing in *B. anynana* followed the protocol in (Prakash & Monteiro 2018). Briefly, Cas9 mRNA was obtained by *in vitro* transcription of linearized pT3TS-nCas9n plasmid (a gift from Wenbiao Chen (Addgene plasmid #46757)) using the mMESSAGE mMACHINE T3 kit (Ambion) and tailed using the Poly(A) Tailing Kit (Ambion) following the manufacturer’s protocol. Guide RNA targets were manually designed by looking for GGN_18_NGG sequence in *dsx* exons, preferably targeting a protein domain. sgRNA templates were prepared according to (Bassett *et al.* 2013) and purified templates were *in vitro* transcribed using T7 RNA polymerase (Roche). 900 ng/µl of purified Cas9 mRNA and 400 ng/µl of purified guide RNA were mixed along with blue food dye and injected into eggs within 2 hours of egg laying. Injections were done with a Borosil glass capillary (World Precision Instruments, 1B100F-3) using a Picospritzer II (Parker Hannifin). Hatched caterpillars were reared on fresh corn leaves and emerging adults were scored for their phenotypes (Supplementary Table 5, 6). Mutated individuals were tested for indels at the targeted site by genomic DNA extraction from thoracic tissues, amplification of targeted regions, cloning and sequencing. All primers and guide RNA sequences are listed in Supplementary Table 4.

### Sexing adults

A W-microsatellite PCR-based method was used to verify the sex of the crispants (van’t Hof *et al.* 2005). The primer sequences that amplify a 185 bp product present only on the W chromosome, were kindly provided by Arjen Van’t Hof and are listed in Supplementary Table 4. PCR was carried out in a 12.5µl reaction volume using 2x PCRBIO Taq Red Mix (PCR Biosystems) and the following protocol: 3 min at 95°C, 40 cycles of 30s at 95°C, 30s at 60°C and 45s at 70°C. Only females produce a band while males do not.

### Genitalia dissections

The genitalia from wildtype and crispants were dissected using fine forceps and placed in 10% solution of sodium hydroxide for 30 min to 1 hour to soften the attached non-sclerotized tissues. They were then moved to PBS and the sclerotized structures were separated from the underlying tissues using forceps. Genitalia were embedded in low melting agarose to maintain orientation while imaging. Imaging was done using an Ocellus microscope (Dun Inc.) consisting of a Canon 7D Mk II DSLR body, Mitutoyo objectives and a P-51 Camlift stacking rail. Individual image slices were processed with Lightroom (Adobe Inc.), stacking of images with Zerene Stacker and post-processing with Photoshop CS5 (Adobe Inc.).

## Acknowledgements and Funding

We thank Arjen van’t Hof for providing the primer sequences to sex *B. anynana* individuals and the canonical male and female *dsx* isoform sequences in *B. anynana*. The partial sequences of the sex-specific *dsx* isoforms generated are available on GenBank (accession nos. MK869725 and MK869726). We also thank Dr. John Ascher and Chui Shao Xiong for help with imaging the dissected genitalia, Zohara Rafi for help in catching *Orsotriaena medus* and Jeffrey Oliver for discussions on phylogenetic hypothesis testing. This work was supported by the Ministry of Education, Singapore (MOE2015-T2-2-159 to A.M), the Department of Biological Sciences, National University of Singapore (R-154-000-608-651 LHK Fund to A.M) and the National University of Singapore (Research Scholarship to A.P).

