## Supplementary Files for "*Doublesex* mediates the development of sex-specific pheromone organs in *Bicyclus* butterflies via multiple mechanisms"

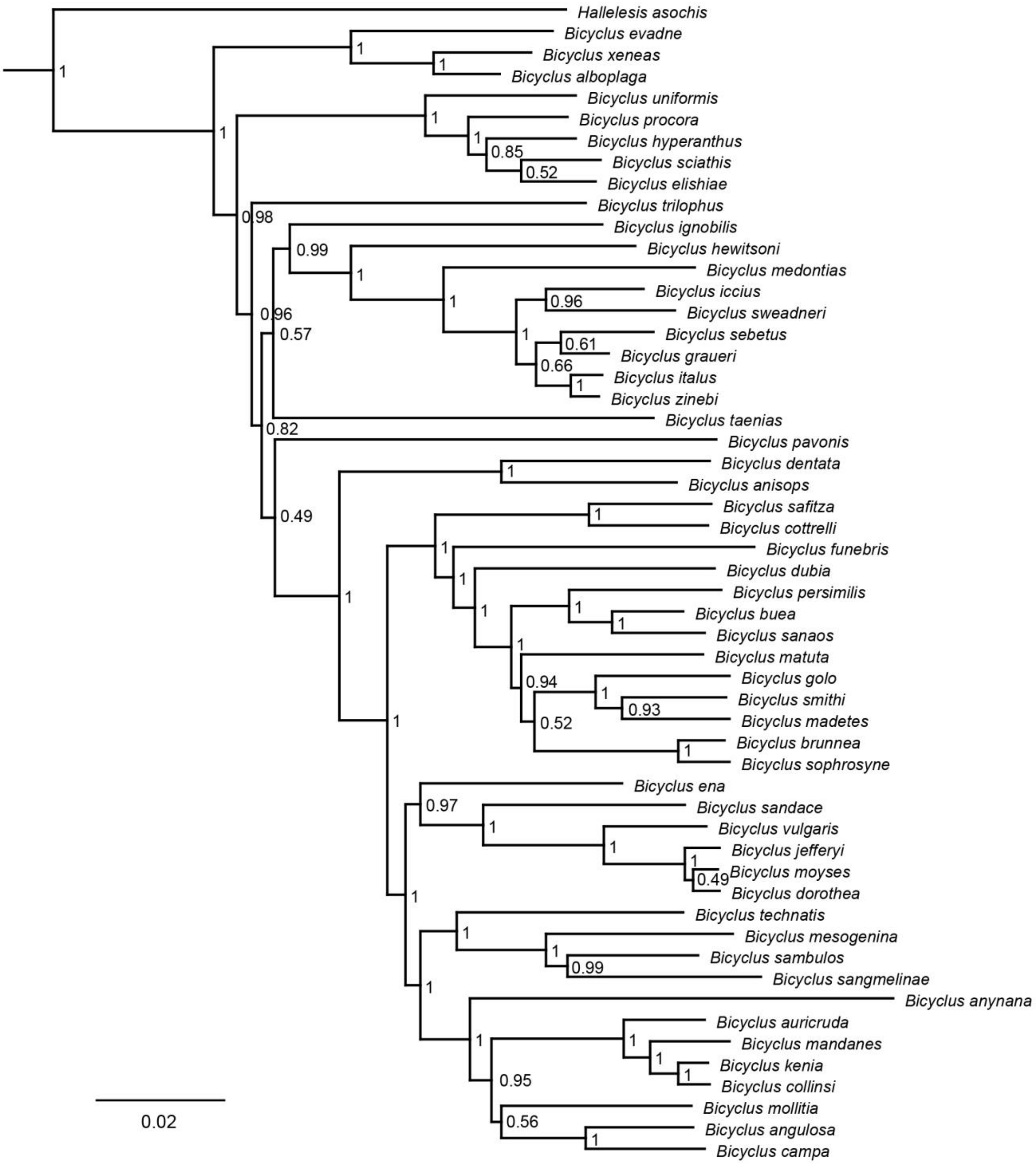


**Supplementary Figure 1: Majority-rule consensus bayesian phylogenetic tree of 53 species of the *Bicyclus* genus with *Hallelesis* as the outgroup.** The tree was generated in MrBayes v3.2.7a. Numbers at the node indicate posterior probabilities.

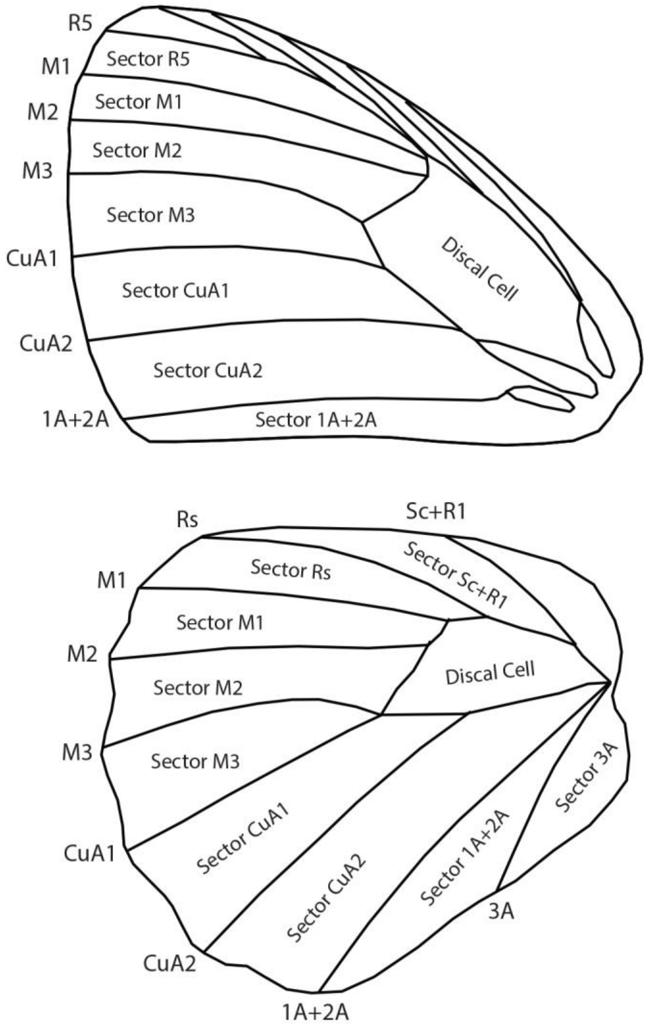


**Supplementary Figure 2: Notation of wing sectors and veins on *Bicyclus anynana* wings.** The names of the sectors (within the wing) and veins (outside the wing) used to reference the different locations on the wing disc is shown.

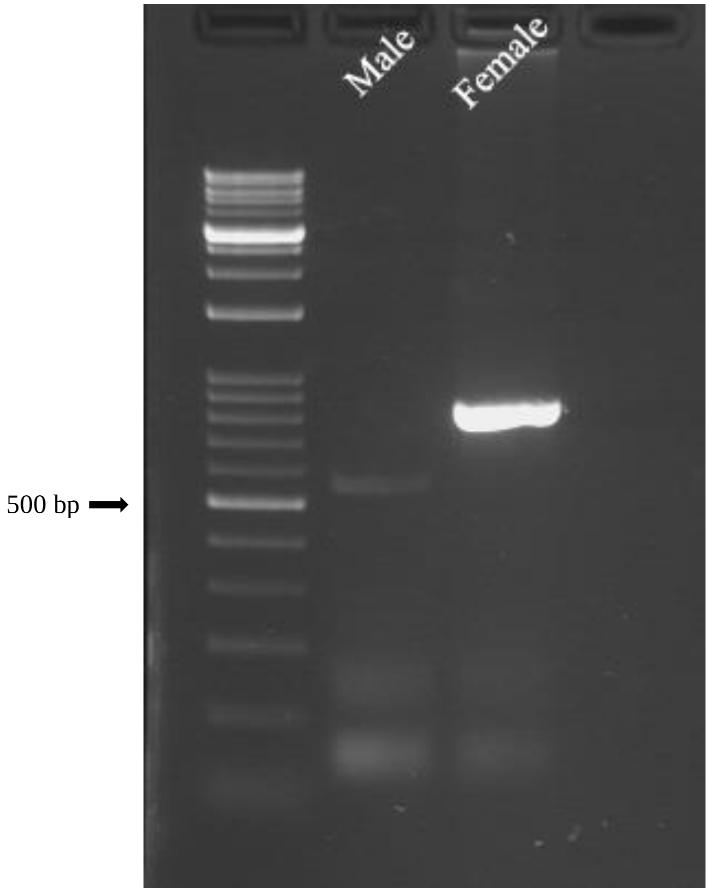


**Supplementary Figure 3: PCR amplification of *dsx* from ~18 hr pupal wing tissues of both sexes of *Bicyclus anynana*.** The wing tissues of each sex express sex-specific isoforms of *dsx*. The partial sequences generated are available on GenBank (accession nos. MK869725 and MK869726).

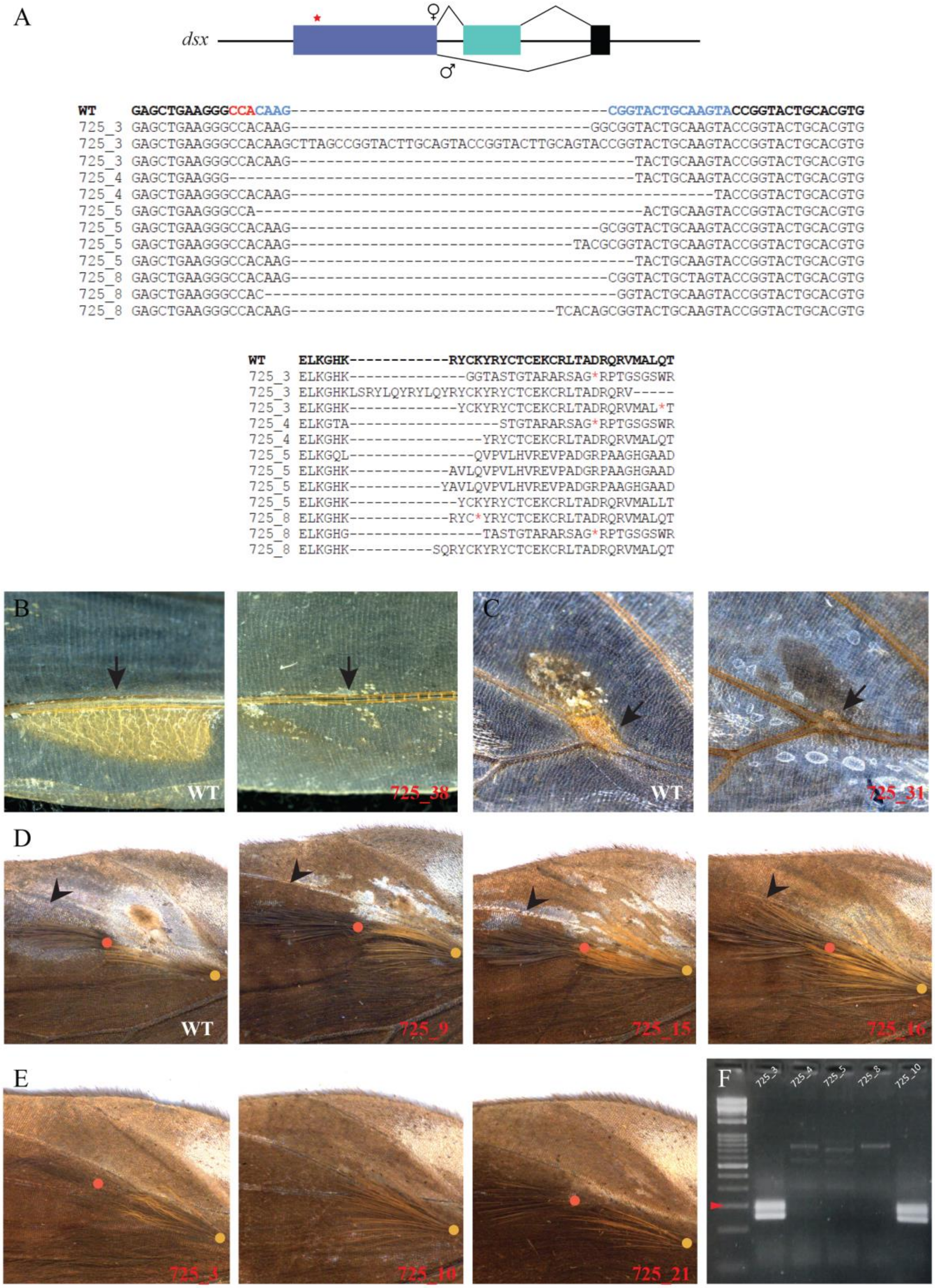
A
dsx
♀
♂
WT GAGCTGAAGGGCCACAAG------------------------------------CGGTACTGCAAGTACCGGTACTGCACGTG
725_3 GAGCTGAAGGGCCACAAG----------------------------------GGCGGTACTGCAAGTACCGGTACTGCACGTG
725_3 GAGCTGAAGGGCCACAAGCTTAGCCGGTACTTGCAGTACCGGTACTTGCAGTACCGGTACTGCAAGTACCGGTACTGCACGTG
725_3 GAGCTGAAGGGCCACAAG---------------------------------------TACTGCAAGTACCGGTACTGCACGTG
725_4 GAGCTGAAGGG----------------------------------------------TACTGCAAGTACCGGTACTGCACGTG
725_4 GAGCTGAAGGGCCACAAG------------------------------------------------TACCGGTACTGCACGTG
725_5 GAGCTGAAGGGCCA--------------------------------------------ACTGCAAGTACCGGTACTGCACGTG
725_5 GAGCTGAAGGGCCACAAG-----------------------------------GCGGTACTGCAAGTACCGGTACTGCACGTG
725_5 GAGCTGAAGGGCCACAAG--------------------------------TACGCGGTACTGCAAGTACCGGTACTGCACGTG
725_5 GAGCTGAAGGGCCACAAG---------------------------------------TACTGCAAGTACCGGTACTGCACGTG
725_8 GAGCTGAAGGGCCACAAG------------------------------------CGGTACTGCTAGTACCGGTACTGCACGTG
725_8 GAGCTGAAGGGCCAC----------------------------------------GGTACTGCAAGTACCGGTACTGCACGTG
725_8 GAGCTGAAGGGCCACAAG------------------------------TCACAGCGGTACTGCAAGTACCGGTACTGCACGTG
WT ELKGHK------------RYCKYRYCTCEKCRLTADRQRVMALQT
725_3 ELKGHK--------------GGTASTGTARARSAG*RPTGSGSWR
725_3 ELKGHKLSRYLQYRYLQYRYCKYRYCTCEKCRLTADRQRV-----
725_3 ELKGHK-------------YCKYRYCTCEKCRLTADRQRVMAL*T
725_4 ELKGTA------------------STGTARARSAG*RPTGSGSWR
725_4 ELKGHK----------------YRYCTCEKCRLTADRQRVMALQT
725_5 ELKGQL---------------QVPVLHVREVPADGRPAAGHGAAD
725_5 ELKGHK------------AVLQVPVLHVREVPADGRPAAGHGAAD
725_5 ELKGHK-----------YAVLQVPVLHVREVPADGRPAAGHGAAD
725_5 ELKGHK-------------YCKYRYCTCEKCRLTADRQRVMALLT
725_8 ELKGHK------------RYC*YRYCTCEKCRLTADRQRVMALQT
725_8 ELKGHG----------------TASTGTARARSAG*RPTGSGSWR
725_8 ELKGHK----------SQRYCKYRYCTCEKCRLTADRQRVMALQT
B
WT
725_38
C
WT
725_31
D
WT
725_9
725_15
725_16
E
725_3
725_10
725_21
F
725_3
725_4
725_5
725_8
725_10

**Supplementary Figure 4: CRISPR/Cas9 mediated *dsx* crispant phenotypes and verification.** A) Top: Schematic of *dsx* male and female isoforms with the region targeted by CRISPR/Cas9 marked with a red star. Bottom: Nucleotide sequences and their translated amino acid sequences from the targeted region of crispant individuals compared with the wildtype sequence in bold. Blue is the target sequence and the PAM sequence is in red. Red stars in the translated protein sequences indicate truncated proteins due to stop codons. (B,C) Loss of patch-associated glands in crispants (black arrows) in comparison to wildtype (WT) D) Hindwing androconia of crispant individuals (in red) compared to WT. Black arrowheads indicate affected development of the greyish-silver scales of patch 2 that lie beneath the black hair-pencil with respect to WT. E) Mutant hair-pencil phenotypes on female hindwings. The yellow and orange dots in (D) and (E) mark the base of hair-pencils as per Figure 1. F) Amplification of female-specific W-microsatellite (~200bp, red arrowhead) to verify the sex of the mutants. Lane 1 is the DNA ladder. Samples 725_3 and 725_10 are females, the others are male.

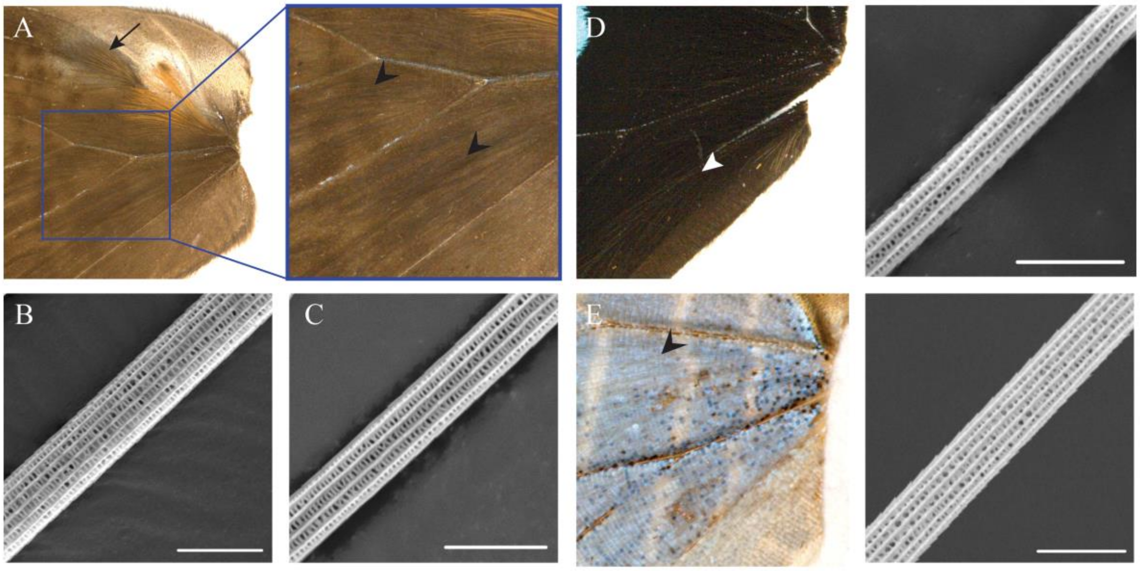


**Supplementary Figure 5: Optical and scanning electron micrographs of hair-pencils and long hairs.** A) Dorsal hindwing of *B. anynana*. Boxed region is expanded to show the long hair-like scales (black arrowheads) that occur on the wings. Black arrow points to the black hair-pencil. B) Scanning electron micrograph (SEM) of a black hair-pencil of *B. anynana*. C) SEM of a long hair-like scale of *B. anynana*. D) Dorsal hindwing of *Papilio palinurus* (left) and the SEM of a long hair-like scale (right), indicated by the white arrowhead. E) Dorsal hindwing of a Lycaenid butterfly (left) and the SEM of a long hair-like scale of this species (right), shown by the black arrowhead. Scale bars: 10µm.

**Supplementary Table 2: Comparison of a single-origin vs multiple-origins hypothesis for different forewing and hindwing hair-pencils and patches among *Bicyclus* species**. The MRCA (Most Recent Common Ancestor) of all lineages that bear the trait of interest was fossilized to either 1 (trait present, single origin) or 0 (trait absent, multiple-origins). Log marginal likelihoods of the two models were calculated using a ReverseJump MCMC analysis on the 15000 post burn-in trees generated from MrBayes. 2*(Δ log marginal likelihood) is the Log Bayes Factors statistic for model testing and values >2 provide positive support towards the better model, which is highlighted in bold.

| **Trait of interest** | **-log marginal likelihood of MCRA = 0** | **-log marginal likelihood of MCRA = 1** | **2*(Δ log marginal likelihood)** |
|---|---|---|---|
| HW Hair-pencil 1 | 10.7185 | **8.66153** | **4.114** |
| HW Hair-pencil 2 | **29.6508** | 30.7024 | **2.1032** |
| HW Hair-pencil 3 | 26.0791 | 26.6614 | 1.165 |
| HW Hair-pencil 4 | **25.2632** | 26.6401 | **2.7538** |
| HW Hair-pencil 6 | **11.3398** | 13.297 | **3.914** |
| FW Hair-pencil 10 | 14.4906 | **12.2945** | **4.392** |
| HW Patch 2 | 30.5968 | 31.4624 | 1.7312 |
| HW Patch 3 | **24.3597** | 27.3627 | **6.006** |
| HW Patch 4 | **24.4543** | 26.8985 | **4.8884** |
| FW Patch 8 | **11.6251** | 14.2916 | **5.333** |
| FW Patch 10 | 26.8476 | 27.7528 | 1.8104 |
| FW Patch 11 | **12.0818** | 13.7735 | **3.38** |
| FW Patch 14 | **15.1644** | 17.2407 | **4.1526** |

| | | | |
|---|---|---|---|
| FW Patch 15 | 31.3586 | 30.7791 | 1.159 |
| FW Patch 16 | 32.6581 | 32.9647 | 0.6132 |
| FW Patch 17 | 11.8391 | **9.5876** | **4.503** |

**Supplementary Table 3: Comparison of a dependent vs independent model of evolution for different pairs of hair-pencils and patches.** Log marginal likelihoods of the two models were calculated using a ReverseJump MCMC analysis on the 15000 post burn-in trees generated from MrBayes. 2*(log marginal likelihood(dependent) – log marginal likelihood (independent)) is the statistic for model testing and values >2 provide support for the dependent model. Pairs of hair-pencils and patches that show correlated evolution are highlighted in bold.

| **Trait 1** | **Trait 2** | **-log marginal likelihood (Dependent)** | **-log marginal likelihood (Independent)** | **2*(Δ log marginal likelihood)** |
|---|---|---|---|---|
| **HW Hair-pencil 1** | **HW Patch 1** | 11.5414 | 15.3335 | **7.60** |
| **HW Hair-pencil 2** | **HW Patch 2** | 46.5223 | 58.1527 | **23.26** |
| **HW Hair-pencil 3** | **HW Patch 3** | 38.8432 | 48.8338 | **19.98** |
| **HW Hair-pencil 4** | **HW Patch 4** | 37.1683 | 47.9479 | **21.56** |
| **HW Hair-pencil 1** | **FW Patch 15** | 37.9627 | 39.4948 | **3.064** |
| HW Hair-pencil 1 | FW Patch 16 | 41.7061 | 41.2099 | -0.9924 |
| HW Hair-pencil 2 | FW Patch 15 | 59.9011 | 58.7345 | -2.33 |
| **HW Hair-pencil 2** | **FW Patch 16** | 51.5141 | 60.3556 | **17.683** |

### Supplementary Table 4: Primers and guide RNA sequences used in this study

| Gene | Primer Number | Primer Sequence |
|---|---|---|
| *dsx* isoform sequencing | AM 27<br>AM 1118 | Forward 5’ ACTGCACGTGCGAGAAGTG 3’<br>Reverse 5’ GAGCAGCACACGCCGTAC 3’ |
| *dsx* common CRISPR Guide | AM 725 | 5’GAAATTAATACGACTCACTATAGGTACTTGCAGTACC GCTTGGTTTTAGAGCTAGAAATAGC 3’ |
| *dsx* common Genotyping | AM 695<br>AM 696 | Forward 5’ GCAATCATCAGTCGCATCGTG 3’<br>Reverse 5’ CAAGTGTGGGAACAGCTAAAG 3’ |
| W-microsatellite primers for sexing *Bicyclus anynana* | AM 797<br>AM 798 | Forward 5’ GAACCAATGCGACAAATGCGACAT 3’<br>Reverse 5’ TGATCTGTATAATCTAATCATAGTGGGTAT AACTAAAACT 3’ |

### Supplementary Table 5: CRISPR/Cas9 injection concentrations and mutation frequencies

| Guide | Guide RNA Conc (ng/µl) | Cas9 mRNA Conc (ng/µl) | Eggs injected | Eggs hatched | Hatch ratio | Total adults | Mutant phenotypes |
|---|---|---|---|---|---|---|---|
| *dsx* common | 400 | 900 | 420 | 166 | 39.52 % | 83 | 32 (38.55%) |

**Supplementary Table 6: Quantification of male and female crispants showing different types of *dsx* crispant phenotypes.**

| Type of *dsx* mutant | Number of males | Number of females |
|---|---|---|
| **Appearance of hair-pencils** | | 13 |
| **Reduction in length of white band** | | 10 |
| **Loss of ventral forewing androconia** | 19 | |
| **Loss of dorsal hindwing androconia** | 18 | |
| **Visible reduction in hair-pencil number** | 5 | |
